# The buoyant cell density of *Cryptococcus neoformans* is affected by capsule size

**DOI:** 10.1101/429936

**Authors:** Raghav Vij, Radames J.B. Cordero, Arturo Casadevall

## Abstract

*Cryptococcus neoformans* is an environmental pathogenic fungus with a worldwide geographical distribution that is responsible for hundreds of thousands human cryptococcosis cases each year. During infection, the yeast undergoes a morphological transformation involving capsular enlargement that increases microbial volume. To understand the factors that play a role in environmental dispersal of *C. neoformans* and *C. gatii* we evaluated the buoyant cell density of *Cryptococcus* using Percoll isopycnic gradients. We found differences in the buoyant cell density of strains belonging to *C. neoformans* and *C. gatti* species complexes. The buoyant cell density of *C. neoformans* strains varied depending on growth medium conditions. In minimal medium, the cryptococcal capsule made a major contribution to the buoyant cell density such that cells with larger capsules had lower density than those with smaller capsules. Removing the capsule, both by chemical or mechanical methods, decreased the *C. neoformans* cell density. Melanization of the *C. neoformans* cell wall, which also contributes to virulence, produced a small but consistent increase in cell density. *C. neoformans* sedimented much slower in seawater as its density approached the density of water. Our results suggest a new function for the capsule whereby it can function as a flotation device to facilitate transport and dispersion in aqueous fluids.

## IMPORTANCE

The buoyant cell density of a microbial cell is an important physical characteristic that may affect its transportability in fluids and interactions with tissues during infection. The polysaccharide capsule surrounding *C. neoformans* is required for infection and dissemination in the host. Our results indicate that the capsule has a significant effect on reducing cryptococcal cell density altering its sedimentation in seawater. Modulation of microbial cell density via encapsulation may facilitate dispersal for other important encapsulated pathogens.

## INTRODUCTION

*C. neoformans* and *gattii* species complexes are important fungal pathogens that can cause pulmonary and serious meningeal disease in humans (1). In the environment, *C. neoformans* is commonly found in soil associated with pigeon excreta, while *C. gattii* is most commonly found on trees (2, 3). *C. gattii* have been isolated from marine and fresh water environments (4, 5). Cryptococcal infection occurs via the respiratory tract where yeast particulates can colonize the lungs (6, 7). In immunocompromised patients, *C. neoformans* can readily disseminate from the lungs to other parts of the body, including the central nervous system by crossing the blood brain barrier. The dissemination of *C. neoformans* yeast cells from the lung to the brain is critical in the development of meningeal disease. The yeast cells undergo drastic morphological changes-during this transition that aid its distribution and evasion from host immune mechanisms. For instance, yeast dimensions can range from 1 to 100 µm in diameter by increasing their cell body and/or growing a thick polysaccharide capsule at the cell wall surface in response to immediate environmental conditions (6–9)(8–11).

The polysaccharide capsule is mostly composed of water (12). It is formed by a porous matrix of branched heteropolysaccharides, mainly glucuronoxylomannan, that extends radially from the cell wall (13). Capsule synthesis is induced under certain stressful conditions, and provides protection against host defense mechanisms by acting as a physical barrier, interfering with phagocytosis and sequestering Reactive Oxygen Species (ROS) and drugs (14, 15). The capsule is essential for the virulence of *C. neoformans* and if of interest for both therapeutic and diagnostic strategies (16).

Melanin is another important virulence factor, such that strains that lack the ability to melanize are less pathogenic (16). Melanin is formed by the polymerization of aromatic and/or phenolic compounds including L-DOPA, methyl-DOPA, epinephrine or norepinephrine (17). In the presence of catecholamine precursors found in the human brain, *Cryptococcus* melanizes its inner cell wall (18). Melanized *C. neoformans* cells are found in the environment (19) and during mammalian infection (20), suggesting an important role of the pigment in *C. neoformans* biology and pathogenesis. Melanization protects cells against a variety of host immune mechanisms and antifungal drugs, as well as, against radiation, desiccation, ROS, and temperature stress (21, 22).

Both the polysaccharide capsule and melanin are complex structures difficult to study. Consequently, it is important to apply biophysical methodologies to gain new insights into the physicochemical properties and biological functions of these major virulence factors (23). One such property that has not been studied in cryptococcal biology is cellular density, presumably a highly-regulated characteristic that may reflect the physiological state of the cell under different conditions (24).

In the first century B.C., Roman writer Vitruvius describes a “Eureka” moment that the Greek polymath Archimedes had when, allegedly, he observed the displacement of water as he sat in a bathtub, which led him to establish the law of buoyancy (25, 26). In a biological context, Archimedes’ law (law of buoyancy) can be applied to calculate the ratio of the absolute mass and volume of an organism which could determine whether it floats or sinks in a fluid of given density. During centrifugation in a continuous Percoll density gradient, cells equilibrate upon reaching the point at which the gradient’s density matches their own. This allows us to estimate buoyant density of *C. neoformans* and *C. gattii* against bead standards of fixed density.

Buoyant density is used for the separation of cell populations but the factors regulating buoyant cell density in microbiology remain understudied, despite the important role it may play in the migration and dissemination of microbial and mammalian cells in fluids. This could be because the buoyant cell density depends on many biological and physical factors, which are often difficult to disentangle. Earlier studies found that the buoyant cell density was affected by the osmolality of the medium in which the cells are grown (27, 28), the encapsulation of bacteria by polysaccharide capsule (29) and the stage of cell cycle (30). Strains of *Porphyromonas gingivalis* with lower buoyant density were less susceptible to phagocytosis, however this could be the result of the correlation between buoyant density and cell surface hydrophobicity (31). Other studies have also reported a difference in buoyant density amongst different strains of mycobacteria and *Burkholderia* spp. (32, 33). In the context of eukaryotes, *Saccharomyces cerevisiae* buoyant cell density varies at different stages of cell cycle (34), and quiescent *S. cerevisiae* cell populations can be separated out using density gradients in a stationary phase culture of the yeast (35).

The buoyant cell density (also referred to as cell density) of *C. neoformans* and the factors that affect it have not been previously investigated. In this study, we use Percoll isopycnic gradients to study the effect of capsule induction, antibody treatment, and melanization on the buoyant cell density of *C. neoformans*.

## RESULTS

### Comparison of *C. neoformans* and *C. gattii* buoyant cell densities

Cell density varied consistently amongst different serotypes of *C. neoformans* and *C. gattii* species complex strains (**Figures 1A&B**). The buoyant density of replicates showed significant variability when comparing *C. neoformans* serotype A (strain H99) to serotype D (strain ATCC 24067) and serotype AD (strain 92.903). However, the density of *C. gatti* did not significantly vary in comparison to *C. neoformans*. To ascertain whether there was a relationship between the density and cell dimension, we imaged the cells with an India-ink counterstain and calculated both the capsule and cell body radii for *C. neoformans* and *C. gatii*. We observed, a statistically significant difference in the cell body radii of all strains when compared to *C. neoformans* serotype A (strain H99). We also observed that the capsule radii of *C. neoformans* Serotype D and AD, and *C. gattii* VG IIa was significantly different when compared to Serotype A.

**Figure 1:**
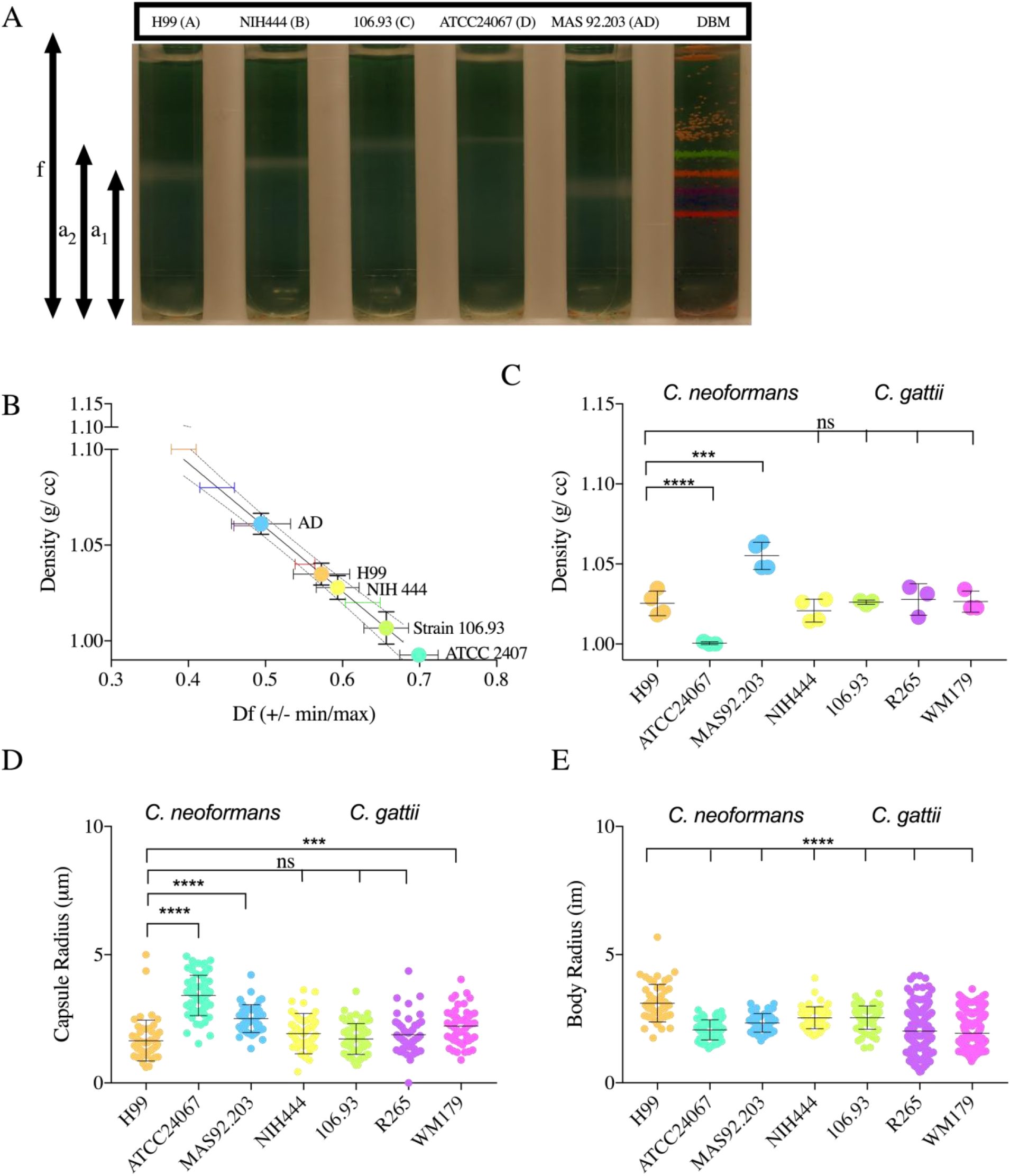
The buoyant cell density of *C. neoformans* and *C. gattii* serotypes. **A.** Representative image of 3–4 independent repetitions of Percoll density gradients comparing the buoyant density of *C. neoformans* (Serotype A, D and AD) and *C. gattii* (Serotype B, C) to density bead markers (DBM). **B.** Representative data from 4 independent experiments depicting the line interpolation of the density factor (min, max) calculated by pixel areas as per the formulae (f – a_1_/f, f – a_2_/f). The df (min, max) values of the density marker beads are used to estimate the buoyant cell density of the cells ran in parallel. **C.** Histogram depicting the difference in buoyant cell density of different serotypes of *C. neoformans* (Serotype A, AD and D) and *C. gattii* (Serotype B, C and variants VGI, VGIIa). The experiment was performed 3 times, as indicated by the symbols on the bar graph, the error bar represents the SD about the mean. **D.** Representative data of capsule *i*. and cell body *ii*. radii of different serotypes and strains. One-way ANOVA was used to for the comparison of cell density, capsule and cell body radii of different strains and serotypes of *C. neoformans* and *C. gattii* to the respective measurements of H99 strain of *C. neoformans*. The following symbols were used to annotate the statistical significance ns (P > 0.05), * P ≤ 0.05), ** (P ≤ 0.01), *** P ≤ 0.001, **** P ≤ 0.0001

### Effect of capsule induction on *C. neoformans* buoyant cell density

In vitro, the capsule is induced in stress conditions such as nutrient starvation medium (36). Cells grown in minimal medium (MM) had significantly lower density (**Figure 2A-C**) in comparison to cells grown in nutrient rich conditions (Sabouraud dextrose broth) where the capsule was significantly smaller. The acapsular strains *cap59* had a significantly higher density than encapsulated cells with the same genetic background. Furthermore, we observed no significant differences in the density of acapsular mutants grown in minimal versus rich medium, confirming the contribution of the polysaccharide capsule in determining the cell density in response to different nutrient conditions.

**Figure 2:**
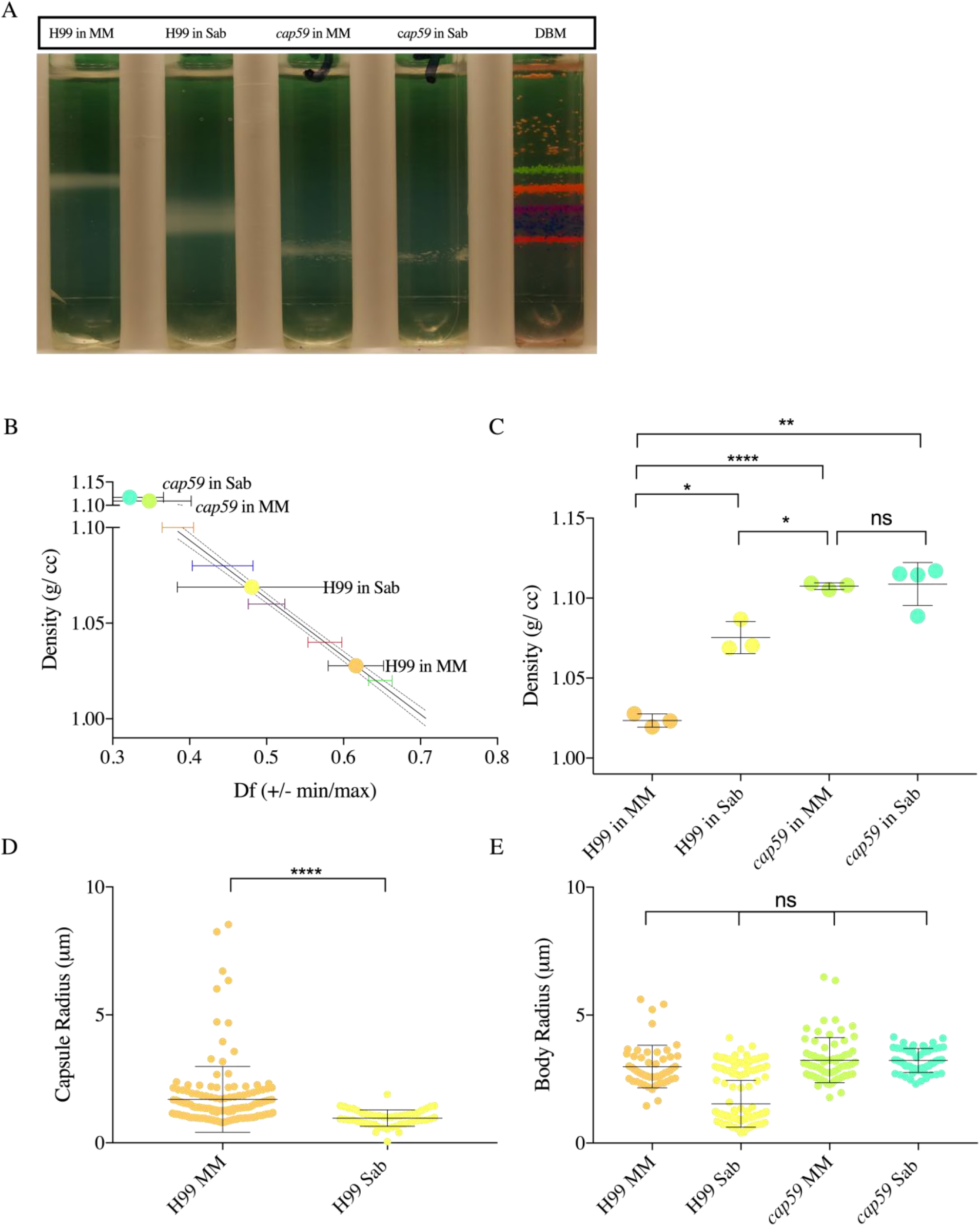
Induction of capsule synthesis decreases *C. neoformans* cell buoyant density. **A.** Representative image of three independent repetitions of Percoll density gradients showing the density of *C. neoformans* H99 juxtaposed with acapsular mutant *Cap59*, both grown in sabouraud medium (Sab), minimal medium (MM). **B.** Representative data from three independent experiments depicting a line interpolation of the density factor with the buoyant densities of the bead standards to calculate the buoyant cell densities of the gradients run in parallel. **C.** Histogram depicting a decrease in the range of buoyant cell density in H99 cells grown in MM, when compared to Sab, due to capsule induction. *Cap59* mutant are significantly denser than normal H99 cells grown in MM. Experiments were performed in replicates independently, as indicated by the data points on the histogram, except for *Cap59* in Sab was performed once as indicated by the symbols on the bar graph, error bar represents SD about the mean. **D.** Representative data depicts *i*. the capsule radii and *ii*. the cell body radii of different strains *C. neoformans* H99 grown in different medium conditions (MM, Sab). *Cap59* grown in MM and Sab do not have a capsule, therefore the capsule radii were not quantified. One-way ANOVA was used to for the comparison of cell density, capsule and cell body radii of *C. neoformans* Cap59 and H99 gown in different conditions. The following symbols were used to annotate the statistical significance ns (P > 0.05), * P ≤ 0.05), ** (P ≤ 0.01), *** P ≤ 0.001, **** P ≤ 0.0001 (For the last two choices only)

Previous studies have reported the molecular composition of the *C. neoformans* capsule by removing the polysaccharide from the cell surface by DMSO extraction and gamma irradiation induced capsule shedding (37). To confirm the effects of the capsule on the buoyant cell density, encapsulated H99 cells were treated with gamma radiation and DMSO to remove capsular material (**Figure 3**). We observed a significant increase in cell density when the capsule was removed by both treatments indicating that the polysaccharide capsule influences the cell density.

**Figure 3:**
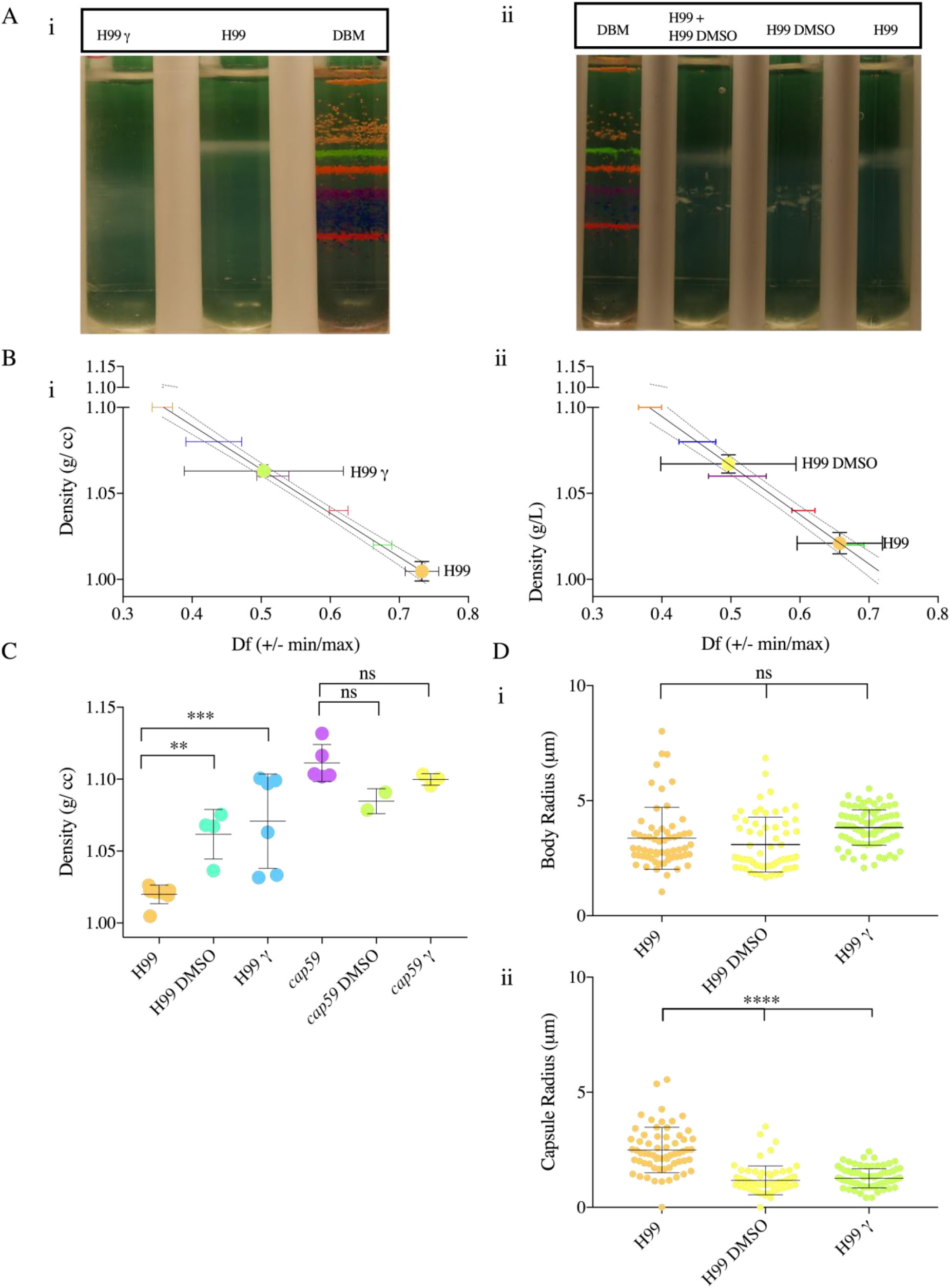
Removal of *C. neoformans* increases the buoyant cell density. **A.** *i* Representative image of four independent repetitions of Percoll density gradients comparing the buoyant cell densities of irradiated () and non-irradiated *C. neoformans* (H99) with a standard of colored uniform density beads. *ii* Representative image of four independent Percoll density gradients of encapsulated *C. neoformans* H99 strains before and after DMSO extraction. **B.** *i, ii* Representative data from independent experiments depicting a line interpolation of the density factor (df) with the buoyant densities of the bead standards, to calculate the buoyant cell densities of *C. neoformans* before and after extraction of the capsule, run in parallel. **C.** A histogram depicting buoyant density of *C. neoformans* before and after capsule extraction by  rays and DMSO. The irradiation experiment was performed independently thrice, and the capsular extraction by DMSO twice, as indicated by the symbols on the bar graph, error bar represents SD about the mean. One-way ANOVA was used to determine differences in density of H99 and Cap59 treated with DMSO and gamma radiation with H99 and Cap59 grown in MM for 10 ds respectively. **D.** Representative data depicts *i*. the capsule radii and *ii*. the cell body radii of *C. neoformans* before and after gamma irradiation capsule shedding. One-way ANOVA was used to for the comparison of cell density, capsule and cell body radii of different strains and serotypes of *C. neoformans* and *C. gattii* to the respective measurements of H99 strain of *C. neoformans*. The following symbols were used to annotate the statistical significance ns (P > 0.05), * P ≤ 0.05), ** (P ≤ 0.01), *** P ≤ 0.001, **** P ≤ 0.0001 (For the last two choices only) interpolation of the density factor with the buoyant densities of the bead standards, to calculate the buoyant cell densities of the gradients run in parallel. **C.** *i*. A histogram depicting density of mel and non-mel *C. neoformans* to compare the density of a mel and non-mel culture started from the same sabouraud pre-culture. The experiment was performed in replicates (n = 4), as indicated by the symbols on the bar graph. A paired t-test found the pairing to be significant (**) and found consistent and significant differences (*) between non melanised and melanised cells. *ii*. A histogram depicting the density of and non-mel cells after removal of capsule by gamma radiation. The experiment was performed in pairs, such that the paired cultures were inoculated from the same sabouraud pre-culture and were treated with gamma-radiation (1500 Gy) together. A paired t-test found the pairing to be significant (*) and found consistent and significant differences (*) between non melanised and melanised cells treated with gamma radiation. **D.** Representative data depicts *i*. the capsule radii and *ii*. the cell body radii of melanized and non-melanized *C. neoformans*. One-way ANOVA was used to for the comparison of capsule and cell body radii of melanised and non-melanised H99 strain of *C. neoformans*. The following symbols were used to annotate the statistical significance ns (P > 0.05), * P ≤ 0.05), ** (P ≤ 0.01), *** P ≤ 0.001, **** P ≤ 0.0001 (For the last two choices only)

### Capsule size correlates with buoyant cell density

Linear regression analysis revealed that capsule radii correlates with cell uch that yeast with larger capsules were less dense (**Figure 4A)**. There was no significant relationship between cell body size and cell density (**Figure 4B**).

**Figure 4:**
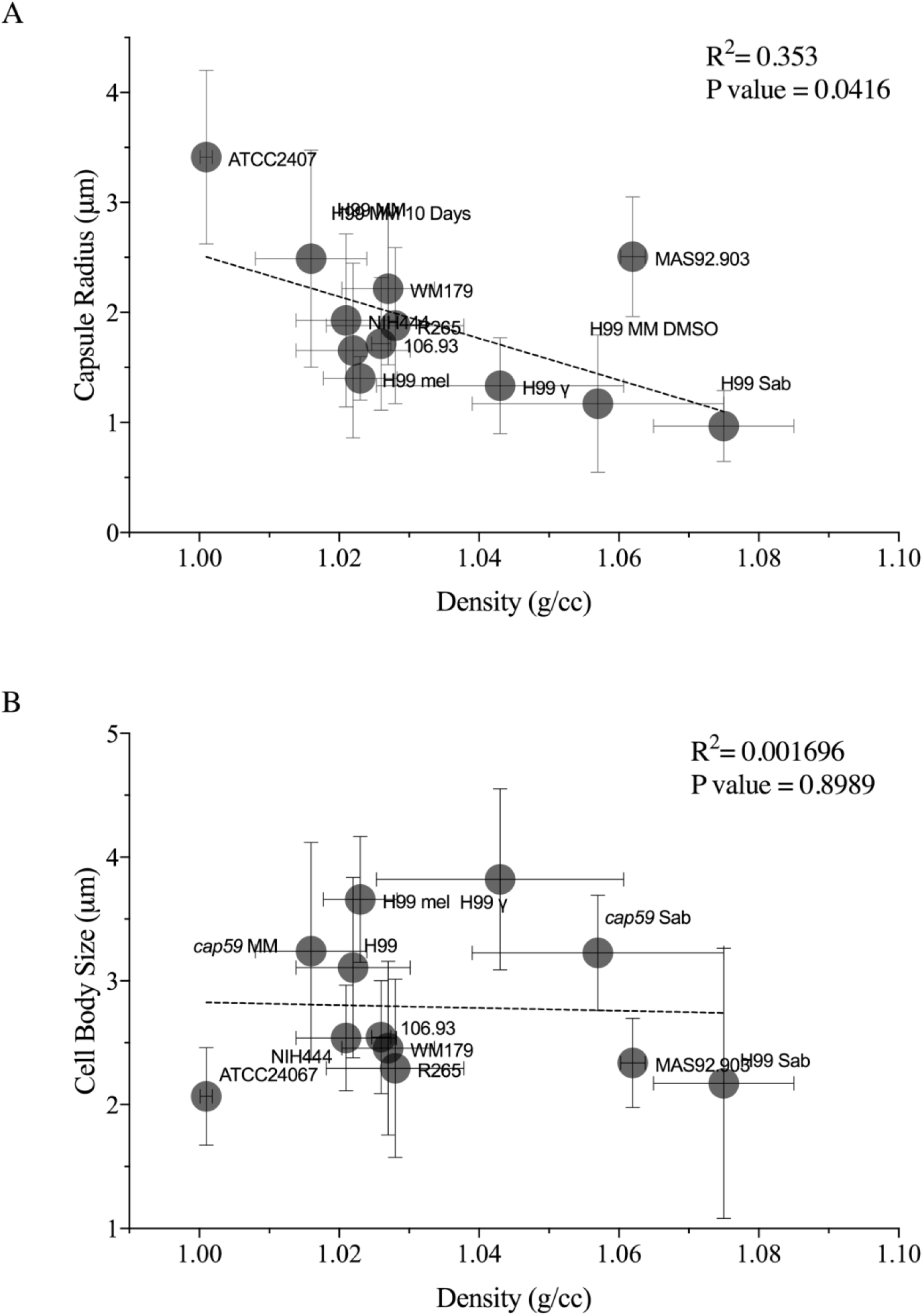
The density of *C. neoformans* and *C. gattii* correlates with the capsule radii. **A.** Density (g/cc) significantly correlates linearly with the capsule size (μm). **B.** No linear relationship was found in when comparing the cell body size (radii) to the density. The density values were collated from all the experiments performed for a specific condition (n = 3 to 5). The cell size was taken from a single experiment for each condition which was found to representative of the replicates.

### Encapsulated *Cryptococcus neoformans* settles slower in sea water

We tested whether the low density of encapsulated *C. neoformans* allowed the fungi to float in water or seawater. The density of seawater is 1.0236 g/cc at room temperature (38, 39), and density of *C. neoformans* in minimal medium is 1.022 +/− 0.008165. When *C. neoformans* grown in minimal medium was added to a cuvette containing seawater, a large population of cells became suspended in the seawater, which became turbid (**Figure 5A**). This effect was not seen with PBS where the cells sink to the bottom within 3 hours (**Figure 5B**).

**Figure 5:**
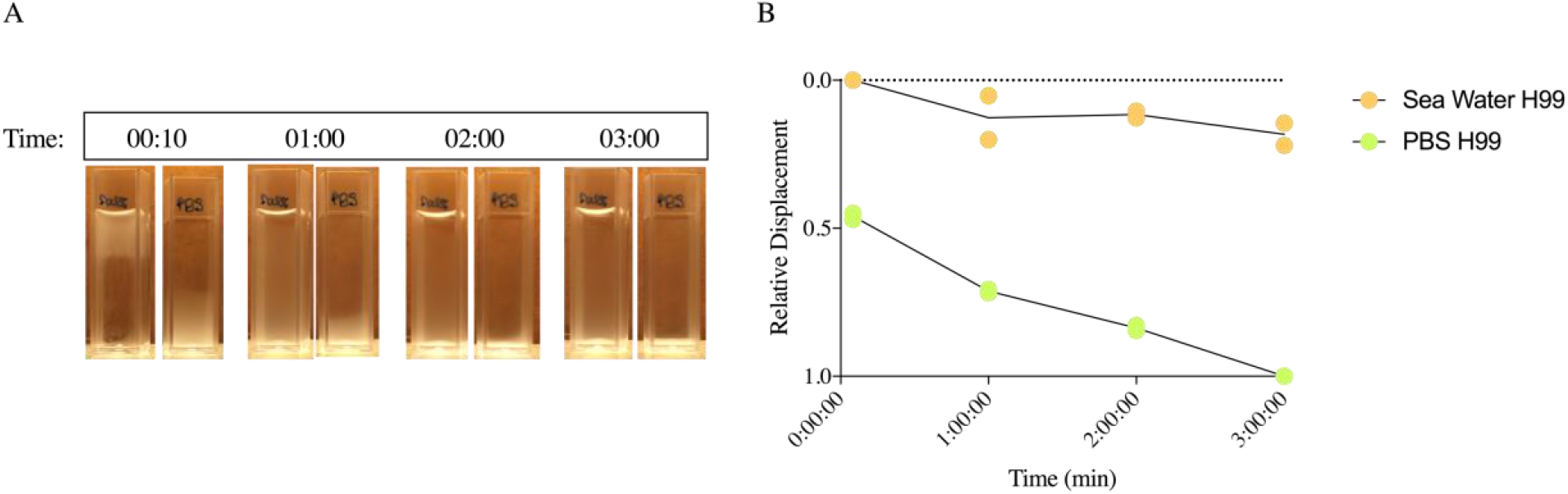
Encapsulated C. neoformans settles slower in seawater. **A.** Representative image of cuvettes (3mL) containing seawater (left) and PBS (right) imaged at different time points shows that H99 grown in MM settles faster in PBS. **B.** Graphical representation of normalized displacement of the upper menisci of cells settling in sea water and PBS. Each data point on a line at a given time-point represents an independent experiment (n=2).

### Melanization increases *C. neoformans* buoyant cell density

Comparison of melanized and non-melanized H99 *C. neoformans* cells demonstrated that melanization was associated with a moderate increase in cell density (**Figure 6)**. Since the increase in the density was small, and melanized cell can easily be distinguished visually from non-melanized cell, we mixed the melanized and non-melanized cells in 1:1 ratio before loading the samples onto the density gradient. While non-melanized cells displayed a range of density that overlapped with melanized cells, the latter tended to have higher density when compared to nonmelanized cells inoculated from the same sabouraud broth pre-culture. Isolated melanin ‘ghosts’(40) had much greater density than cells, estimated to be > 1.1 g/ cc (data not shown). Note that melanized cells also had smaller capsules (**Figure 6D, ii**), which may contribute to the increase in cell density. Thus, we also compared the density of melanized and non-melanized cells after removal of the capsule by gamma radiation (**Figure 6C, ii**). Upon capsule removal, we observed that the density of melanized cells was consistently and significantly higher than non-melanized cells.

**Figure 6:**
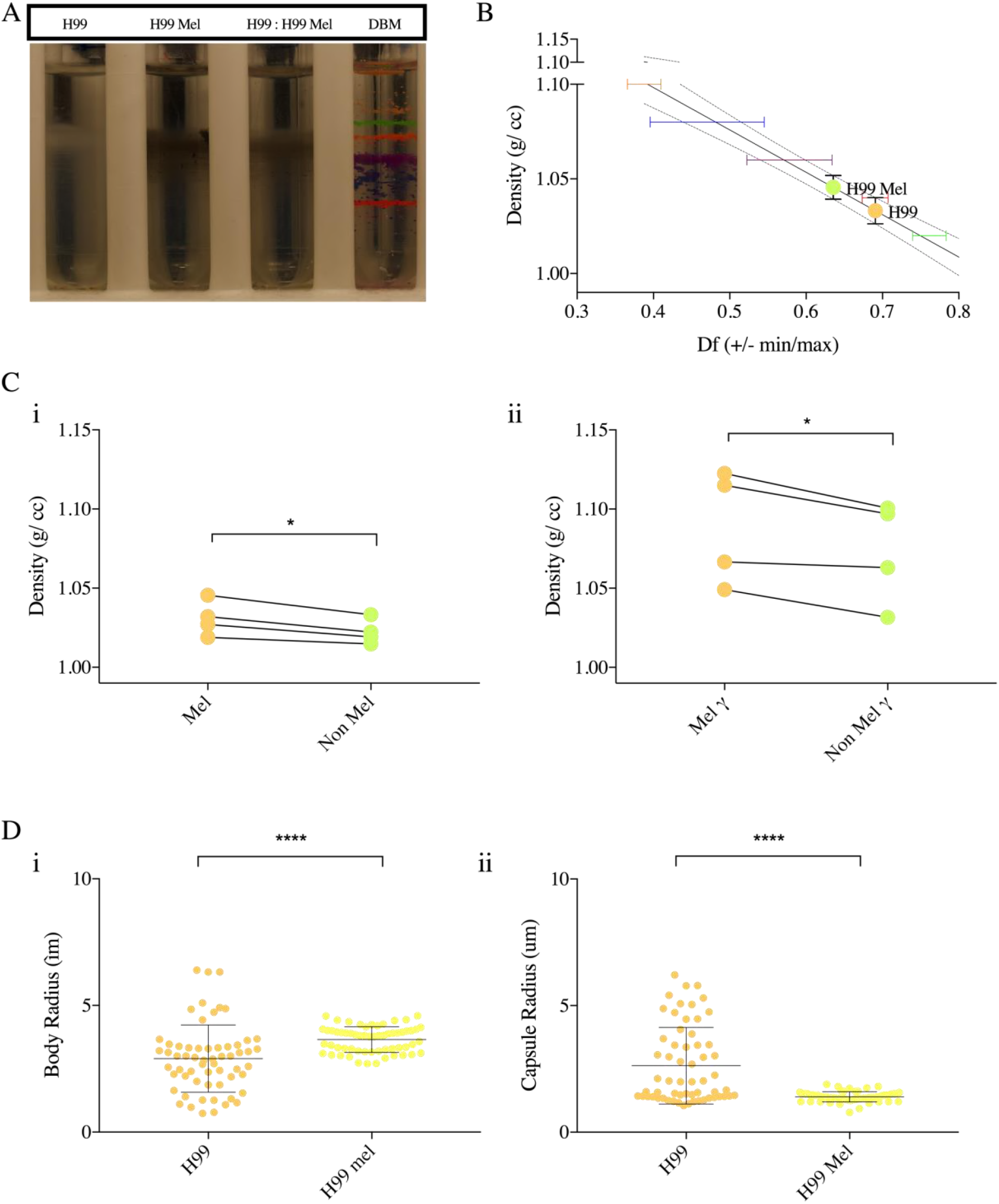
Figure 6: Effect of Melanization on *C. neoformans* buoyant cell density. **A.** Representative image of four independent repetitions of Percoll density gradients comparing the density of H99 in MM, H99 in MM with L-DOPA (mel) and a 1:1 mixture of the cells. The white cells (H99) band slightly above the melanized black cells (mel) as can be seen by the gradient that contains the mixture. **B.** Representative data from three independent experiments depicting a line interpolation of the density factor with the buoyant densities of the bead standards, to calculate the buoyant cell densities of the gradients run in parallel. **C.** *i*. A histogram depicting density of mel and non-mel *C. neoformans* to compare the density of a mel and non-mel culture started from the same sabouraud pre-culture. The experiment was performed in replicates (n=4), as indicated by the symbols on the bar graph. A paired t-test found the pairing to be significant (**) and found consistent and significant differences (*) between non melanised and melanised cells. *ii*. A histogram depicting the density of and non-mel cells after removal of capsule by gamma radiation. The experiment was performed in pairs, such that the paired cultures were inoculated from the same sabouraud pre-culture and were treated with gamma-radiation (1500 Gy) together. A paired t-test found the pairing to be significant (*) and found consistent and significant differences (*) between non melanised and melanised cells treated with gamma radiation. **D.** Representative data depicts *i*. the capsule radii and *ii*. the cell body radii of melanized and non-melanized *C. neoformans*. One-way ANOVA was used to for the comparison of capsule and cell body radii of melanised and non-melanised H99 strain of *C. neoformans*. The following symbols were used to annotate the statistical significance ns (P > 0.05), * P ≤ 0.05), ** (P ≤ 0.01), *** P ≤ 0.001, **** P ≤ 0.0001 (For the last two choices only)

### Antibody binding and other conditions that have no significant effect on *C. neoformans* buoyant cell density

Previous studies have shown that capsular antibodies alter the viscoelastic properties and structure of the capsule (41). Antibody binding also causes a change in the hydration state of the PS capsule (12). Treatment of H99 *C. neoformans* with capsular antibodies (18B7 and E1) did not significantly alter the cell buoyant density (Figure **S1**). Furthermore, binding of mouse complement, incubations at different salt concentration (to induce osmotic stress) and incubation in lipid rich medium had no significant impact on *C. neoformans* buoyant cell density (Figure **S2**).

**Figure S1:**
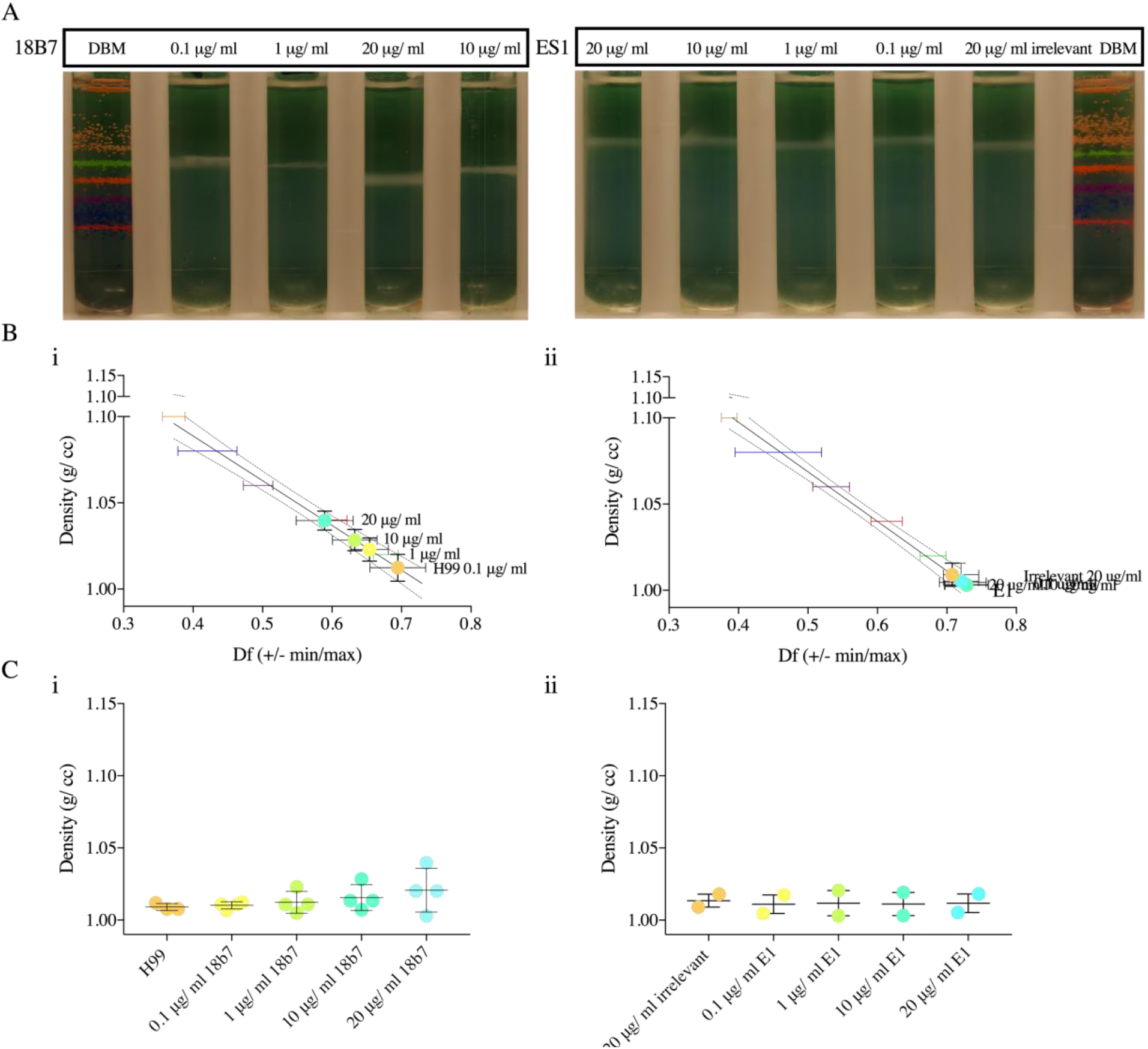
Binding of capsular antibodies do not alter *C. neoformans* buoyant density. **A.** Representative image of independent repetitions of Percoll density gradients comparing the density of *C. neoformans* H99 with and without antibody incubation with i. 18B7 and ii. ES1. **B.** Representative data independent experiments depicting a line interpolation of the density factor with the buoyant densities of the bead standards, to calculate the buoyant cell densities of the gradients run in parallel. **C.** *i*. A histogram depicting density of *C. neoformans* H99 incubated with capsular antibodies i. 18B7 (n=4) and ii. ES1 (n=2) at different concentrations (0.1,1,10,20 μg/mL)

**Figure S2.**
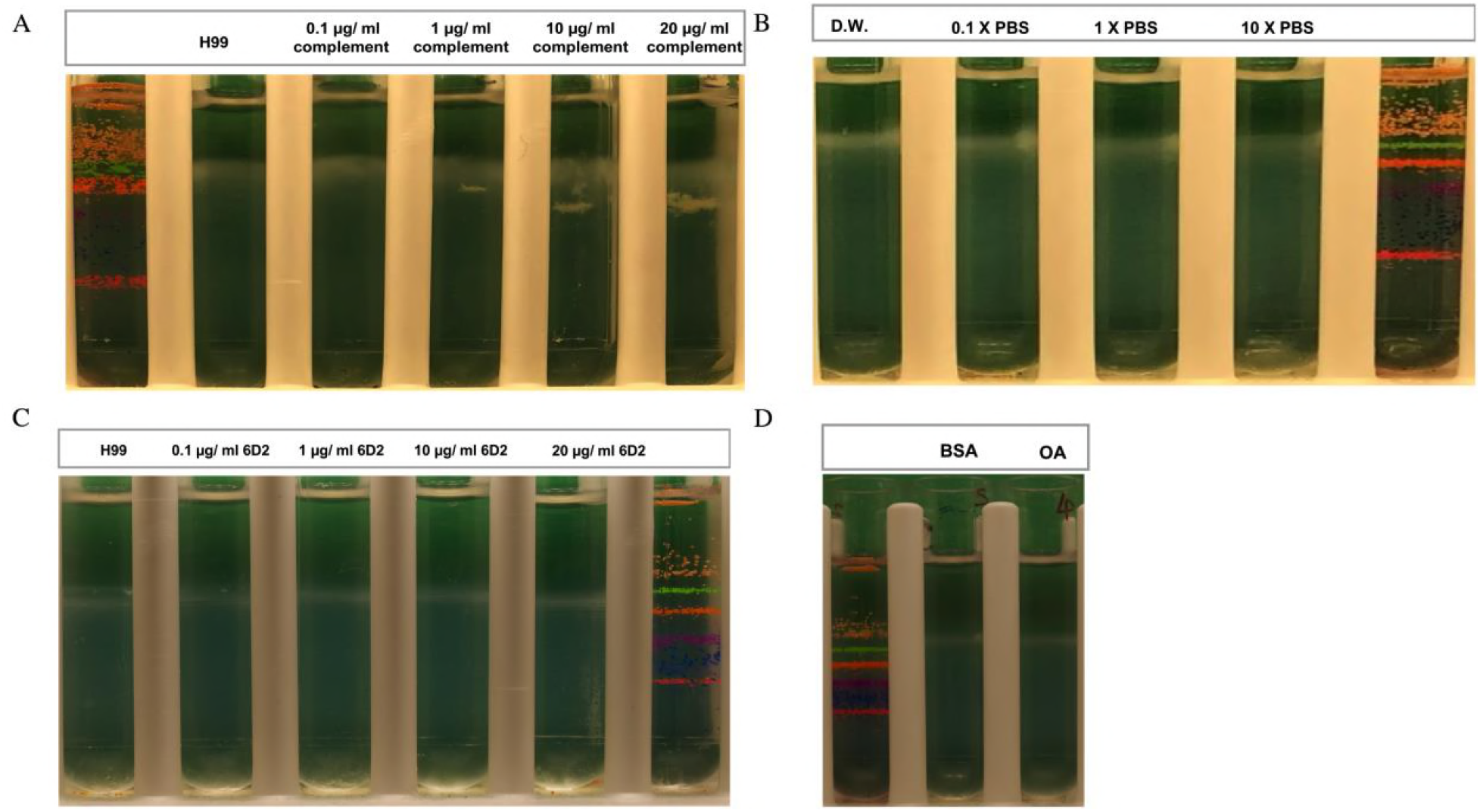
Effect of complement binding, osmotic shock, melanin-binding antibody and growth in rich lipid media on buoyant density. Image of Percoll density gradients comparing the density of *C. neoformans* H99 upon **A.** complement binding, **B.** osmotic shock, **C.** 6D2 antibody binding **D.** growth in lipid rich media. These conditions do not affect the buoyant cell density of *Cryptococcus neoformans* (H99) significantly. Experiments were done once.

## DISCUSSION

In this study, we characterized the buoyant cell density of *C. neoformans* and *C. gattii* in different conditions. We report minor differences in buoyant densities between serotypes of the cryptococcal species complex. Our results also suggest that the capsule plays a major role in determining the buoyant cell density of the yeast such that encapsulated strains have densities close to that of water. Meanwhile, melanization increased the density slightly. Changes in buoyancy could influence the dispersal of the yeast in the environment, and dissemination of the fungal pathogen during infection. Furthermore, the buoyant density may be used for the separation of different populations of yeast cells (35), to characterize *C. neoformans* mutants with capsular defects (16) and for the isolation of the titan cells (42).

The buoyant density of a microbe is a fundamental biophysical property that influences its behavior in aqueous fluids. Depending on its density, a microbe could remain suspended in a fluid or settle to the bottom. Amongst other factors, this could influence the microbe’s access to nutrients, sunlight and oxygen. Thus, it is not surprising that marine and freshwater unicellular organisms including phytoplankton, regulate their cell density via mechanisms that involve the synthesis and storage of gas vacuoles, polysaccharide mucilage sheaths, and glycogen (43). Interestingly, the polysaccharide mucilage sheath of these bacteria, that resembles the polysaccharide capsule of *C. neoformans*, has been characterized as an important factor that decreases the density of the cell to just below the density of water (44). Our data demonstrates that the cryptococcal capsule serves a similar function by increasing the volume of the yeast cell without significantly increasing its mass and thereby reducing its density.

A quantitative parameter used to determine how fast a population of microbial cells sinks in a fluid of given density is the settling velocity, which is calculated by the Stoke’s law and depends on the buoyant density and the size (diameter) for a spherical object such as a yeast cell (43, 45). In marine bacteria, low cell density (< 1.064 g/cc) correlates with low settling velocity as calculated by Stoke’s law (46). The variable size of *C. neoformans* grown in minimal medium (3–16 μm) and the low density (∼ 1.022 g/cc) we observed during nutrient starvation conditions in an aqueous environment suggests that the settling velocity of *C. neoformans* would be similarly low. More importantly, the encapsulated *C. neoformans* cells would have a lower settling velocity when compared to similar-sized cells that have no capsule due to the decrease in density.

We hypothesize that the capsule can function as a flotation device that allows the yeast cell to flow horizontally in aqueous environment to access (47) nutrients, oxygen and disperse the pathogen (48). For instance, a study found that a *C. gattii* clinical isolate survived in filtered ocean water, distilled water and saline water (up-to 10% of initial inoculum) at room temperature up to 94 days (49). The resistance of *Cryptococcus* to different levels of osmotic stress is consistent with our observations that high salt concentrations do not alter the cell density. The strains of *C. neoformans* and *C. gattii* have heterogeneous global distribution, and the mechanism of the dispersal are unknown (50). Possibly; the varied density of the strains influences the differential dispersal of the fungal pathogen. Thus, in the context of environmental fungal pathogens *C. neoformans* and *gattii*, the cell density could play an important role in determining the dispersal of the yeast in the environment and affect its ability to infect a wide range of hosts, including marine mammals such as dolphins (51–53). Estimating the settling velocity of microbial cells in aqueous fluids will add weight to the hypothesis that the buoyant density of *C. neoformans* and *C. gattii* influences environmental dispersal.

Melanization had a moderate influence on cell density. Despite the much greater density of melanin ghosts, cellular melanization had a small effect on cell density presumably due to the fact that melanin contributes approximately only 15.4% (m/m) of cellular mass (40).

In immunocompromised hosts, the *C. neoformans* can disseminate from the lungs to the brain where it causes life-threatening meningitis. This multistep process could require the fungal cell to travel into the draining lymph node and into fluidic blood and lymph systems to survive and grow outside the lungs. Murine models have shown that the capsule and cell body size is different at different sites of infection (9). Our results showing that the capsule makes an important contribution to reducing buoyant density and increasing flotation suggests that the capsular enlargement that occurs during infection could be a major variable in determining dissemination in body fluids.

In summary, the density of *C. neoformans* grown in minimal medium is slightly greater than that of water. The presence of a capsule reduced the density such that it approached that of water. Hence, the capsule, by reducing density, also reduces the settling velocity of *C. neoformans* in aqueous solutions, which could favor environmental dispersal. The establishment of *C. gattii* in the Pacific Northwest is reported to have occurred relatively recently (50). Although the means by which *C. gattii* reached North America are unknown the fact that it has been recovered from marine environments (49) together with our finding of a low density of yeast cells indicating a propensity of the pathogen to float suggest that sea currents could have transported *C. gattii* between continents. The observation that the polysaccharide capsule makes a large contribution to reducing density suggests a new role for this structure in the environment as an aid to cell dispersal and transport in aqueous fluids.

## Acknowledgement

AC was supported by grants 5R01HL059842, 5R01AI033774, 5R37AI033142, and 5R01AI052733. We would like to thank Dr. Francoise Dromer (Institut Pasteur) for providing capsular antibody ES1 (IgG). R.V. designed and conducted the experiments, analyzed the data and wrote the manuscript. R.J.B.C. and A.C. contributed to the experimental design, supervised the experiments, edited and wrote parts of the manuscript.

## Materials and Methods

### Yeast cultures

Frozen stocks of *C. neoformans* and *gatti* strains were inoculated into Sabouraud agar rich medium (pH adjusted to 7.4) at 30°C for 48 hours. Yeast cultures of *C. neoformans* included

| Species | Strain | Reference or Source |
| --- | --- | --- |
| *C. neoformans* | H99 | John Perfect (Durham, NC) |
|  | Cap59 | (54) |
| *C. neoformans* | ATCC24067 | ATCC (Manassas, VA) |
| *C. neoformans hybrids* | MAS92-203 | (55) |
| *C. gatii* | NIH444, ATCC32609 | (55) |
|  | 106.93 | (55) |
|  | VGI, WM179 | ATCC (Manassas, VA) |
|  | VGIIa, R265 | ATCC (Manassas, VA) |

 Acapsular mutants from *Cryptococcus neoformans* cultured included Cap59 (Background H99, serotype A). Approximately 10^6^ cells from the stationary phase cultures in Sabouraud broth were washed twice in Minimal Medium (10 mM MgSO4, 29.3 mM KH2PO4, 13 mM glycine, 3 µM thiamine-HCl, and 15 mM dextrose with pH adjusted to 5.5). The washed cells were inoculated into Minimal Medium (MM) for capsule induction, MM with L-DOPA (100 mM) to induce melanization, and Sabouraud broth for providing rich medium conditions. Cells were incubated at 37°C for 48 hours, rotating at 180 RPM. Cells were washed twice with sterile PBS (Phosphate Buffer Saline), centrifuging them for 5 minutes at 4700 x *g*. Cells were counted using a hemocytometer, and dilutions were made to obtain 1 X 10^7^ cells in PBS. The cells were then loaded onto Percoll Density gradients with or without treatments to test the effect of different conditions on the buoyant cell density.

### Density gradient centrifugation

Percoll is a non-toxic and isotonic alternative to the commonly used sucrose gradient, and is composed of polyvinylpyrrolidone coated colloidal silica particles (56). Percoll has found applications for separation of mammalian blood, tumor, immune and endothelial cells, and microbial cells due to its ability to form reproducible self-generated continuous gradients (57). Stock Isotonic Percoll (SIP) was obtained by added 1 part of 1.5 M NaCl to 9 parts of Percoll. The working solution of 70% (v/v) was obtained by diluting SIP with 0.15 M NaCl, to a final density of 1.0914 g/ml. Three milliliters of this solution were loaded into polycarbonate ultracentrifuge tubes (13 X 51 mm). Approximately, 10^7^ cells were pelleted at 4700 x *g* and over layered directly or after treatment. All gradients were run in parallel with a standard tube.

For the preparation of the standard tube, 10 µl of each uniform density bead standard (Cospheric DMB kit) including light orange (ORGPMS-1.00 250–300um, density 1.00 g/cc), fluorescent green (1.02 g/cc), florescent orange (1.04 g/cc), florescent violet (1.06 g/cc), dark blue (1.08) and florescent red (1.099 g/cc), was loaded and mixed with the Percoll.

By varying time and speed of centrifugation, it was found that the most optimal separation of the density gradient beads, which was taken as an indication for the most optimal continuous density gradient formed, occurred at 40,000 RPM for 30 minutes (acceleration 9, deceleration 0), in TLA 100.3 fixed angle rotor in Optima TLX tabletop ultracentrifuge.

### Buoyant cell density estimation

First, the images of the density gradient were taken under uniform light and shadow conditions using Nikon D3000 DSLR, Auto settings. Next, pixel area measurements were taken from the bottom of the tube, to the area at the beginning of each band (a1), ranging to the end of each band (a2), to the upper meniscus of the tube (f). The density factor, *Df (min, max)*, and the average along with the standard deviation was computed on Microsoft Xcel according to the following formulae,

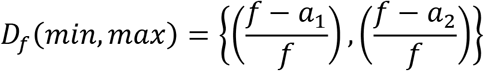

A standard curve was derived, where *Df* (min, max) and buoyant density (g/l) of the density marker beads were computed by linear regression. A 95% confidence interval was used to interpolate the mean density of sample cells, around a standard deviation, run in parallel with the uniform density bead standards.

Although, the results from different Percoll gradient runs follow the same trend, the exact density values can vary considerably. This can be attributed to pipetting errors or errors in measurement of density factor.

### Gamma irradiation of cells for capsule removal

Gamma irradiation was used to remove the capsule as described earlier (58). Approximately 10^9^ cells of melanized and non-melanized cells were plated on a 24-well plate. The cells were irradiated to a total dose of 1500 Gy, using Shepherd Mark 1 at the SKCCC Experimental Irradiator Core at Johns Hopkins University Sidney Kimmel Comprehensive Cancer Center. Cells were washed twice in PBS and approximately 10^7^ cells were pelleted at 4700 x *g* and loaded onto the gradient.

### DMSO Extraction of *C. neoformans* capsule

Approximately 10^7^ cells were incubated in 15 ml of DMSO at 30°C for 30 minutes to allow capsule extraction. The cells were washed thrice in 1X PBS, pelleted and loaded onto the Percoll density gradient.

### Antibody Coating of *C. neoformans* capsule

Purified antibodies, 18B7 and E1 (kindly provided by the Dromer’s laboratory), were obtained from stock solutions kept at 4°C. The antibodies were serially diluted in PBS to concentrations of 20µg, 10µg, 1 µgand 0.1 µg/ml. A pellet of 10^7^ cells was suspended with 1 ml of each Ab solution in Eppendorf tubes, vortexed and incubated at 28°C on a rotating mixer, for 1 hour.

### *C. neoformans* melanization

Frozen stalks of *C. neoformans* H99 was inoculated into Sabouraud broth and incubated at 30°C for 48 hours, till the cultures reached stationary phase. The cells were counted using a hemocytometer. 10^6 cells/ml of were inoculated into minimal medium (10 mM MgSO4, 29.3 mM KH2PO4, 13 mM glycine, 3 µM thiamine-HCl, and 15 mM dextrose with pH adjusted to 5.5) with and without L-DOPA (100 mM). The cells were cultured for 10 days at 30°C rotating at 180 RPM. The cells were washed twice in PBS, and 10^7 cells were pelleted at 47000 x *g* loaded of melanized, non-melanized and 1:1 mixture of the cells was loaded onto the gradient. Melanin ghosts were prepared as described (40).

### Mouse complement deposition in *C. neoformans*

Frozen stocks of guinea pig complement (1 mg/ml) were thawed. 50% (500 g/ml), 20%, 10% and 1% dilutions with PBS was added a pellet of 10^7^ cells. The cells were incubated with complement for 1 hour at 28°C in a rotating-mixer.

### Providing *C. neoformans* with osmotic stress

*C. neoformans* (approximately 10^8^ cells) were incubated with 1 ml of 10X PBS, 1X PBS, 0.1X PBS and ultrapure distilled water (MilliQ) for 2 hours in 30 rotating-mixer at 28°C.

### Settling of *C. neoformans* in seawater

10^7^ cells of C. neoformans grown in MM were gently pipetted onto cuvettes containing 3ml of sea water (Worldwide Imports AWW84130 Live Nutri Seawater) and PBS. The settling of the cells was observed by imaging the cuvettes at different time intervals with a Nikon D3000 DSLR. The images were analyzed using image J. The relative displacement was measured using the following formulae (f-u)/f, where f is the area of the tube, u is the area from the bottom of the tube to the upper menisci of cells that are settling.

### Cell imaging and yeast size measurements

The cells were visualized and imaged with India Ink negative staining under Olympus AX70 Microscope at 20X magnification and 40X magnification. The capsule and cell body size was estimated using an automated measurement Python software (59) or by ImageJ when cells were observed to be aggregated.

### Statistical analysis

All statistical analysis was performed on GraphPad Prism 7.0 software. The density of cells was estimated by using making a standard curve from beads of different densities using linear regression to estimate the unknown values of a given sample with a 95% confidence interval. Details of statistical tests applied are provided on the figure legends.

## Reference

1. Perfect JR. 2000. Cryptococcosis, p. 79–93. In Atlas of Infectious Diseases. Current Medicine Group, London.

2. Ellis DH, Pfeiffer TJ. 1990. Natural habitat of Cryptococcus neoformans var. gattii. J Clin Microbiol 28:1642–1644.

3. Emmons CW. 1960. Prevalence of Cryptococcus neoformans in pigeon habitats. Public Health Rep 75:362–364.

4. Bartlett KH, Kidd SE, Kronstad JW. 2008. The emergence of Cryptococcus gattii in British Columbia and the Pacific Northwest. Curr Infect Dis Rep 10:58–65.

5. MacDougall L, Kidd SE, Galanis E, Mak S, Leslie MJ, Cieslak PR, Kronstad JW, Morshed MG, Bartlett KH. 2007. Spread of Cryptococcus gattii in British Columbia, Canada, and Detection in the Pacific Northwest, USA. Emerg Infect Dis 13:42–50.

6. Neilson JB, Fromtling RA, Bulmer GS. 1977. Cryptococcus neoformans: size range of infectious particles from aerosolized soil. Infect Immun 17:634–638.

7. Powell KE, Dahl BA, Weeks RJ, Tosh FE. 1972. Airborne Cryptococcus neoformans: particles from pigeon excreta compatible with alveolar deposition. J Infect Dis 125:412–415.

8. Feldmesser M, Kress Y, Casadevall A. 2001. Dynamic changes in the morphology of Cryptococcus neoformans during murine pulmonary infection. Microbiology 147:2355–2365.

9. Charlier C, Chrétien F, Baudrimont M, Mordelet E, Lortholary O, Dromer F. 2005. Capsule Structure Changes Associated with Cryptococcus neoformans Crossing of the Blood-Brain Barrier. Am J Pathol 166:421–432.

10. Zaragoza O, García-Rodas R, Nosanchuk JD, Cuenca-Estrella M, Rodríguez-Tudela JL, Casadevall A. 2010. Fungal Cell Gigantism during Mammalian Infection. PLoS Pathog 6.

11. Okagaki LH, Strain AK, Nielsen JN, Charlier C, Baltes NJ, Chrétien F, Heitman J, Dromer F, Nielsen K. 2010. Cryptococcal Cell Morphology Affects Host Cell Interactions and Pathogenicity. PLOS Pathog 6:e1000953.

12. Maxson ME, Cook E, Casadevall A, Zaragoza O. 2007. The volume and hydration of the Cryptococcus neoformans polysaccharide capsule. Fungal Genet Biol 44:180–186.

13. Cordero RJB, Pontes B, Guimarães AJ, Martinez LR, Rivera J, Fries BC, Nimrichter L, Rodrigues ML, Viana NB, Casadevall A. 2011. Chronological Aging Is Associated with Biophysical and Chemical Changes in the Capsule of Cryptococcus neoformans. Infect Immun 79:4990–5000.

14. Casadevall A, Coelho C, Cordero RJB, Dragotakes Q, Jung E, Vij R, Wear MP. 2018. The Capsule of Cryptococcus neoformans. Virulence 0.

15. Zaragoza O, Rodrigues ML, De Jesus M, Frases S, Dadachova E, Casadevall A. 2009. The capsule of the fungal pathogen Cryptococcus neoformans. Adv Appl Microbiol 68:133–216.

16. Kwon-Chung KJ, Rhodes JC. 1986. Encapsulation and melanin formation as indicators of virulence in Cryptococcus neoformans. Infect Immun 51:218–223.

17. Garcia-Rivera J, Eisenman HC, Nosanchuk JD, Aisen P, Zaragoza O, Moadel T, Dadachova E, Casadevall A. 2005. Comparative analysis of Cryptococcus neoformans acid-resistant particles generated from pigmented cells grown in different laccase substrates. Fungal Genet Biol 42:989–998.

18. Nosanchuk JD, Stark RE, Casadevall A. 2015. Fungal Melanin: What do We Know About Structure? Front Microbiol 6.

19. Nosanchuk JD, Rudolph J, Rosas AL, Casadevall A. 1999. Evidence That Cryptococcus neoformans Is Melanized in Pigeon Excreta: Implications for Pathogenesis. Infect Immun 67:5477–5479.

20. Nosanchuk JD, Rosas AL, Lee SC, Casadevall A. 2000. Melanisation of Cryptococcus neoformans in human brain tissue. The Lancet 355:2049–2050.

21. Casadevall A, Rosas AL, Nosanchuk JD. 2000. Melanin and virulence in Cryptococcus neoformans. Curr Opin Microbiol 3:354–358.

22. Cordero RJB, Casadevall A. 2017. Functions of fungal melanin beyond virulence. Fungal Biol Rev 31:99–112

23. Frases S, Viana NB, Casadevall A. 2011. Biophysical Methods for the Study of Microbial Surfaces. Front Microbiol 2.

24. Grover WH, Bryan AK, Diez-Silva M, Suresh S, Higgins JM, Manalis SR. 2011. Measuring single-cell density. Proc Natl Acad Sci 108:10992–10996.

25. Biello D, Biello D. 2006. Fact or Fiction?: Archimedes Coined the Term “Eureka!” in the Bath. Sci Am.

26. Kuroki H. 2016. How did Archimedes discover the law of buoyancy by experiment? Front Mech Eng 11:26–32.

27. Baldwin WW, Myer R, Powell N, Anderson E, Koch AL. 1995. Buoyant density ofEscherichia coli is determined solely by the osmolarity of the culture medium. Arch Microbiol 164:155–157.

28. Baldwin WW, Kubitschek HE. 1984. Evidence for osmoregulation of cell growth and buoyant density in Escherichia coli. J Bacteriol 159:393–394.

29. Lowe BA, Miller JD, Neely MN. 2007. Analysis of the Polysaccharide Capsule of the Systemic Pathogen Streptococcus iniae and Its Implications in Virulence. Infect Immun 75:1255–1264.

30. Kubitschek HE. 1987. Buoyant density variation during the cell cycle in microorganisms. Crit Rev Microbiol 14:73–97.

31. Sundqvist G, Figdor D, Hänström L, Sörlin S, Sandström G. 1991. Phagocytosis and virulence of different strains of Porphyromonas gingivalis. Eur J Oral Sci 99:117–129.

32. Sagripanti J-L, Carrera M, Robertson J, Levy A, Inglis TJJ. 2011. Size distribution and buoyant density of Burkholderia pseudomallei. Arch Microbiol 193:69–75.

33. Vijay S, Nair RR, Sharan D, Jakkala K, Mukkayyan N, Swaminath S, Pradhan A, Joshi NV, Ajitkumar P. 2017. Mycobacterial Cultures Contain Cell Size and Density Specific Sub-populations of Cells with Significant Differential Susceptibility to Antibiotics, Oxidative and Nitrite Stress. Front Microbiol 8.

34. Baldwin WW, Kubitschek HE. 1984. Buoyant density variation during the cell cycle of Saccharomyces cerevisiae. J Bacteriol 158:701–704.

35. Allen C, Büttner S, Aragon AD, Thomas JA, Meirelles O, Jaetao JE, Benn D, Ruby SW, Veenhuis M, Madeo F, Werner-Washburne M. 2006. Isolation of quiescent and nonquiescent cells from yeast stationary-phase cultures. J Cell Biol 174:89–100.

36. Zaragoza O, Casadevall A. 2004. Experimental modulation of capsule size in Cryptococcus neoformans. Biol Proced Online 6:10–15.

37. Frases S, Nimrichter L, Viana NB, Nakouzi A, Casadevall A. 2008. Cryptococcus neoformans Capsular Polysaccharide and Exopolysaccharide Fractions Manifest Physical, Chemical, and Antigenic Differences. Eukaryot Cell 7:319–327.

38. Nayar KG, Sharqawy MH, Banchik LD, Lienhard V JH. 2016. Thermophysical properties of seawater: A review and new correlations that include pressure dependence. Desalination 390:1–24.

39. Sharqawy MH, Lienhard JH, Zubair SM. 2010. Thermophysical properties of seawater: a review of existing correlations and data. Desalination Water Treat 16:354–380.

40. Wang Y, Aisen P, Casadevall A. 1996. Melanin, melanin “ghosts,” and melanin composition in Cryptococcus neoformans. Infect Immun 64:2420–2424.

41. Cordero RJB, Pontes B, Frases S, Nakouzi AS, Nimrichter L, Rodrigues ML, Viana NB, Casadevall A. 2013. Antibody Binding to Cryptococcus neoformans Impairs Budding by Altering Capsular Mechanical Properties. J Immunol Author Choice 190:317–323.

42. Zaragoza O, Nielsen K. 2013. Titan cells in Cryptococcus neoformans: cells with a giant impact. Curr Opin Microbiol 16:409–413.

43. Reynolds CS, Oliver RL, Walsby AE. 1987. Cyanobacterial dominance: The role of buoyancy regulation in dynamic lake environments. N Z J Mar Freshw Res 21:379–390.

44. Reynolds CS. 2007. Variability in the provision and function of mucilage in phytoplankton: facultative responses to the environment. Hydrobiologia 578:37–45.

45. Richardson TL, Jackson GA. 2007. Small Phytoplankton and Carbon Export from the Surface Ocean. Science 315:838–840.

46. Inoue K, Nishimura M, Nayak BB, Kogure K. 2007. Separation of marine bacteria according to buoyant density by use of the density-dependent cell sorting method. Appl Env Microbiol 73:1049–1053.

47. Condie SA, Bormans M. 1997. The Influence of Density Stratification on Particle Settling, Dispersion and Population Growth. J Theor Biol 187:65–75.

48. Kidd SE, Bach PJ, Hingston AO, Mak S, Chow Y, MacDougall L, Kronstad JW, Bartlett KH. 2007. Cryptococcus gattii Dispersal Mechanisms, British Columbia, Canada. Emerg Infect Dis 13:51–57.

49. Kidd SE, Chow Y, Mak S, Bach PJ, Chen H, Hingston AO, Kronstad JW, Bartlett KH. 2007. Characterization of Environmental Sources of the Human and Animal Pathogen Cryptococcus gattii in British Columbia, Canada, and the Pacific Northwest of the United States. Appl Environ Microbiol 73:1433–1443.

50. Roe CC, Bowers J, Oltean H, DeBess E, Dufresne PJ, McBurney S, Overy DP, Wanke B, Lysen C, Chiller T, Meyer W, Thompson GR, Lockhart SR, Hepp CM, Engelthaler DM. 2018. Dating the Cryptococcus gattii Dispersal to the North American Pacific Northwest. mSphere 3.

51. Gales N, Wallace G, Dickson J. 1985. Pulmonary cryptococcosis in a striped dolphin (Stenella coeruleoalba). J Wildl Dis 21:443–446.

52. Migaki G, Gunnels RD, Casey HW. 1978. Pulmonary cryptococcosis in an Atlantic bottlenosed dolphin (Tursiops truncatus). Lab Anim Sci 28:603–606.

53. Miller WG, Padhye AA, van Bonn W, Jensen E, Brandt ME, Ridgway SH. 2002. Cryptococcosis in a Bottlenose Dolphin (Tursiops truncatus) Caused by Cryptococcus neoformans var. gattii. J Clin Microbiol 40:721–724.

54. Coelho C, Souza ACO, Derengowski L da S, de Leon-Rodriguez C, Wang B, Leon-Rivera R, Bocca AL, Gonçalves T, Casadevall A. 2015. Macrophage mitochondrial and stress response to ingestion of Cryptococcus neoformans. J Immunol Baltim Md 1950 194:2345–2357.

55. Cordero RJB, Frases S, Guimaräes AJ, Rivera J, Casadevall A. 2011. Evidence for branching in cryptococcal capsular polysaccharides and consequences on its biological activity. Mol Microbiol 79:1101–1117.

56. Pertoft H, Rubin K, Kjellén L, Laurent TC, Klingeborn B. 1977. The viability of cells grown or centrifuged in a new density gradient medium, Percoll (TM). Exp Cell Res 110:449–457.

57. Pertoft H. 2000. Fractionation of cells and subcellular particles with Percoll. J Biochem Biophys Methods 44:1–30.

58. Bryan RA, Zaragoza O, Zhang T, Ortiz G, Casadevall A, Dadachova E. 2005. Radiological studies reveal radial differences in the architecture of the polysaccharide capsule of Cryptococcus neoformans. Eukaryot Cell 4:465–475.

59. Dragotakes Q, Casadevall A. 2018. Automated Measurement of Cryptococcal Species Polysaccharide Capsule and Cell Body. J Vis Exp JoVE.

